# Passive acoustic monitoring as a tool for early detection of invasive animal species: a systematic review and a case study

**DOI:** 10.64898/2026.04.24.720520

**Authors:** David Becker, Marit K. Kasten, Torben Weber, Ingo Grass, Thomas Hiller

## Abstract

Invasive animal species are spreading rapidly across the globe, creating an urgent need for efficient early-detection and monitoring tools. Passive acoustic monitoring has become an established method in biodiversity research, but its application to invasive species monitoring has been less systematically explored. Here, we combine a systematic literature review with a field-based case study to evaluate the potential of passive acoustic monitoring for invasive animal detection. We identified 26 studies on acoustic monitoring of invasive animals, mainly addressing amphibians (11 studies), birds and fish (five each) with most studies from the USA and Australia. The use of acoustic monitoring of invasive species has increased during the past decade, with recent studies applying automated detection, machine learning, and large-scale monitoring frameworks. As a case study, we further tested the feasibility of low-cost acoustic monitoring of the invasive American bullfrog (*Lithobates catesbeianus*) in southwestern Germany, combined with automated identification using BirdNET. We successfully confirmed bullfrog presence in eight of the eleven monitored lakes, including sites close to a protected nature reserve. Our results highlight the growing potential of passive acoustic monitoring of invasive species under field conditions. In combination with automated species detection, manual validation, and emerging real-time monitoring devices, passive acoustic monitoring becomes an increasingly powerful tool for early intervention and scalable management of biological invasions.

## Introduction

Globalization and worldwide trade have created unprecedented connectivity between regions and continents, allowing species to be transported far beyond their native ranges. This makes the introduction and establishment of non-native species a common consequence of human activity (Hulme 2009; Early et al. 2016; Seebens et al. 2017; Pyšek et al. 2020). While many introduced species do not cause immediate or noticeable impacts, some can become invasive, resulting in substantial ecological and economic consequences (Simberloff et al. 2013; Pyšek et al. 2020). Invasive organisms can alter community structure, disrupt trophic interactions, facilitate pathogen transmission, and thereby contribute to the decline of native species (Bellard et al. 2016; Blackburn et al. 2019).

Once invasive populations establish and spread, eradication is often technically difficult and costly, making prevention and early monitoring the most cost-effective management strategies (Leung et al. 2002; Hulme 2006; Simberloff et al. 2013; Lodge et al. 2016). However, monitoring rapidly spreading species, particularly across large spatial scales, is resource-intensive and often constrained by limited financial capacity for manpower (Haubrock et al. 2021). Traditional monitoring approaches such as transect walks or trapping include repeated site visits and can therefore be difficult to sustain over time, particularly when species are nocturnal, or only seasonally active (Haubrock et al. 2021).

In this context, novel monitoring technologies are increasingly explored to improve detection efficiency and spatial coverage (Sugai et al. 2019; Gibb et al. 2019). Passive acoustic monitoring, which relies on autonomous recording units to capture soundscapes over extended time periods, has emerged as a promising tool for biodiversity assessment and wildlife monitoring (Shonfield & Bayne 2017; Sugai et al. 2019). Because many animals produce species-specific vocalizations, acoustic monitoring allows detection without continuous human presence and can operate at temporal and spatial scales that exceed traditional field surveys (Sugai et al. 2019; Gibb et al. 2019). Technical advances enable automated species identifications from large datasets of audio recordings (Knight et al. 2017; Kahl et al. 2021). Convolutional neural networks, such as BirdNET, are widely used applications to automatically identify species of varying taxa including birds and amphibians (Kahl et al. 2021).

Although passive acoustic monitoring has already long been applied in biodiversity research (Kalko & Schnitzler 1989, Riede 1993), its targeted application for invasive species monitoring has only recently become more common (Ribeiro et al. 2022; Bota et al. 2024; Wood et al. 2024). Studies across multiple taxa indicate that acoustic methods can contribute substantially to the detection and monitoring of invasive animal populations (Taylor et al. 2017; Roe et al. 2018; Bota et al. 2024; McEwen et al. 2024; Wood et al. 2024; Jenkins et al. 2025; Leung et al. 2025). These studies cover a broad spectrum of applications, including species-specific detection trials as well as larger-scale monitoring implementations (Roe et al. 2018; Leung et al. 2025). However, a comprehensive overview of how passive acoustic monitoring has been used for invasive species across taxa and ecosystems is missing.

Among invasive vertebrates, amphibians are particularly suitable candidates for passive acoustic monitoring and automated species identification due to their distinct mating calls (Gerhardt 1994; Adams & Pearl 2007; Urbina et al. 2020). The American bullfrog (*Lithobates catesbeianus* Shaw, 1802) is one of the most widespread invasive amphibian species worldwide and is posing a serious threat to native biodiversity (Adams & Pearl 2007; Adriaens et al. 2013; Urbina et al. 2020) (Figure 1 C+D). Native to eastern North America, American bullfrogs have established populations in South America, Europe, and Asia through aquaculture, pet trade, and intentional releases (Laufer & Waitzmann 2002, 2020; Adriaens et al. 2013). Their high annual reproductive output, broad diet and tolerance of anthropogenic habitats facilitate rapid population growth and expansion (Nussbaum & Storm 1983; Adams & Pearl 2007; Urbina et al. 2020). In Europe, populations continue to spread locally, raising concerns for native amphibians and wetland ecosystems (Bota et al. 2024).

**Figure 1:**
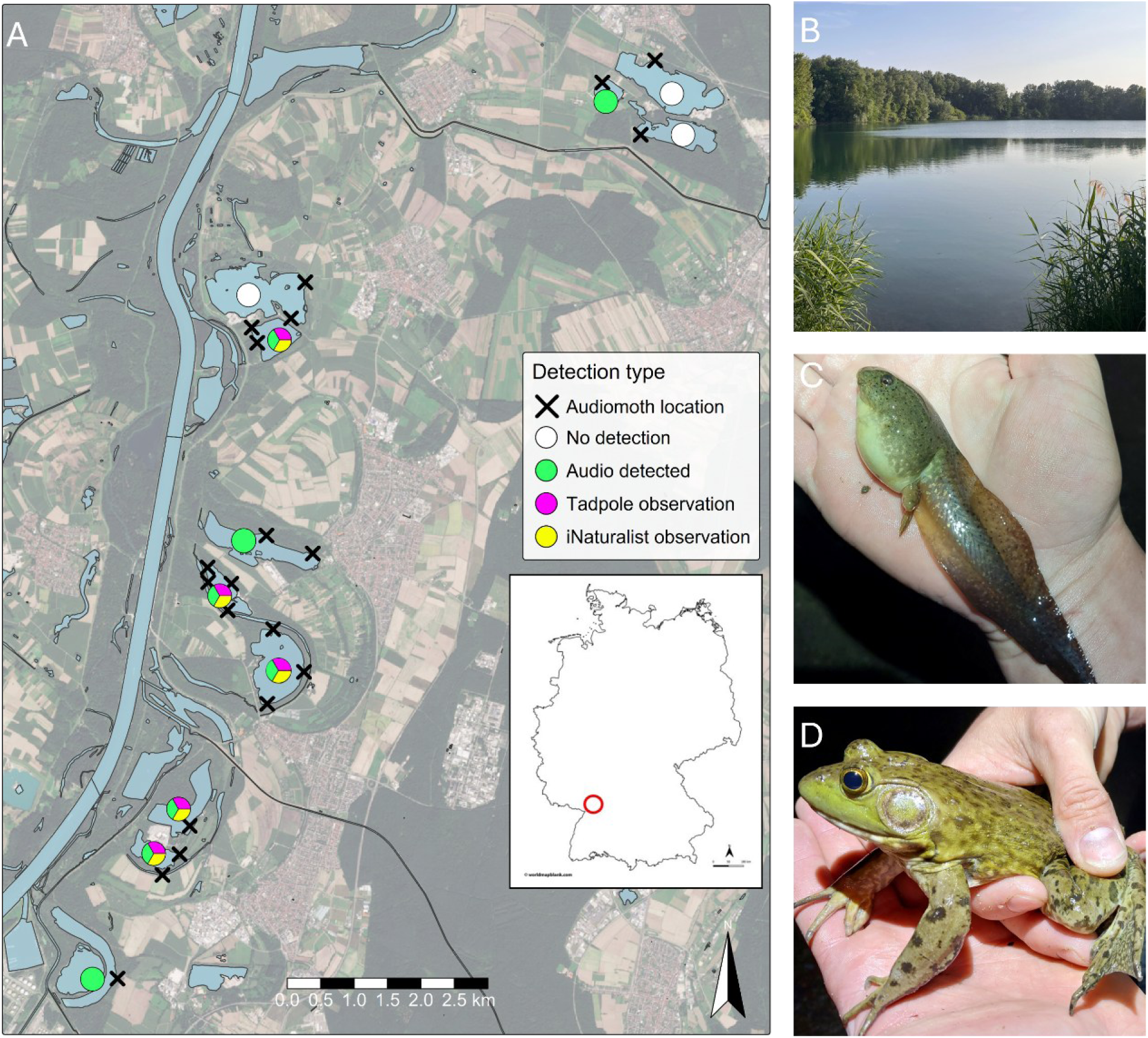
(A) Study lakes in the Upper Rhine region near Karlsruhe, Germany (red circle in inset map), showing AudioMoth deployment locations (black crosses) and bullfrog detections recorded by different methods (coloured circles) **(B)** Panoramic view of Lake Streitköpflesee, supposed source pool of invasive American bullfrogs. **(C)** Tadpole and **(D)** adult American bullfrog (by Lennhard Hudel, CC-BY, iNaturalist observation ID C=242549731; D= 216431325).

In southwestern Germany, American bullfrogs have established expanding populations in gravel pit lakes around the Upper Rhine region since the early 2000s (Baden-Württemberg, near Karlsruhe) (Laufer & Waitzmann 2002), where the nature conservation authorities are trying to limit the negative impacts of further regional spread (Laufer & Waitzmann 2020; Wengerter et al. 2024). The proximity of several protected nature reserves to known bullfrog populations raises concern about the conservation of native species, particularly given the limited capacity for systematic monitoring. Therefore, automatically detecting newly colonized water bodies at an early stage is crucial to assess invasion dynamics and inform management responses. However, monitoring multiple lakes consistently over extended periods of time presents logistical and financial challenges for local authorities.

Given these challenges, this study combines a systematic literature review with a field-based case study to evaluate the potential of passive acoustic monitoring in invasive species monitoring. We synthesize existing applications across taxa to assess current developments and remaining gaps. Furthermore, we tested the feasibility of low-cost acoustic monitoring and automated species detection for invasive American bullfrogs in German lakes. Integrating global evidence with local acoustic data allowed us to examine whether automated acoustic species detection can provide reliable and management-relevant information that could support more efficient and scalable monitoring of invasive species.

## Passive acoustic monitoring of invasive animal species: systematic literature review

### Methods

To identify relevant literature, we used Scopus with the search string TITLE-ABS-KEY((bullfrog* OR “*Lithobates catesbeianus*” OR “invasive species” OR invasive) AND (”acoustic monitoring” OR “bioacoustic*” OR “passive acoustic monitoring”) AND NOT “non-invasive”) on 6.11.2025. This search resulted in 143 papers. We screened their abstracts to narrow down to 44 potentially relevant papers. After assessing the full texts of those 44 papers, we kept 26 relevant papers. We excluded studies that did not directly address the application of passive acoustic monitoring for invasive animal species, post-removal ecosystem restoration, invasive plants, or other monitoring methods such as telemetry.

### Results

The 26 identified papers covered a broad range of invasive animal taxa across terrestrial and aquatic ecosystems worldwide (Figure 2A; Supporting Information Table S1). Amphibians were the most frequently studied group (11 studies), followed by birds and fish (five each), arthropods (three), and mammals and reptiles (one each). Repeated focal species included cane toads (*Rhinella marina*) in Australia and barred owls (*Strix varia*) in the USA, alongside studies on different invasive amphibians, fish, and crayfish in Europe, North and South America, and Asia (Figure 2A). The earliest studies in our dataset were published in 2012, with increasing publication numbers in more recent years (Figure 2B).

**Figure 2:**
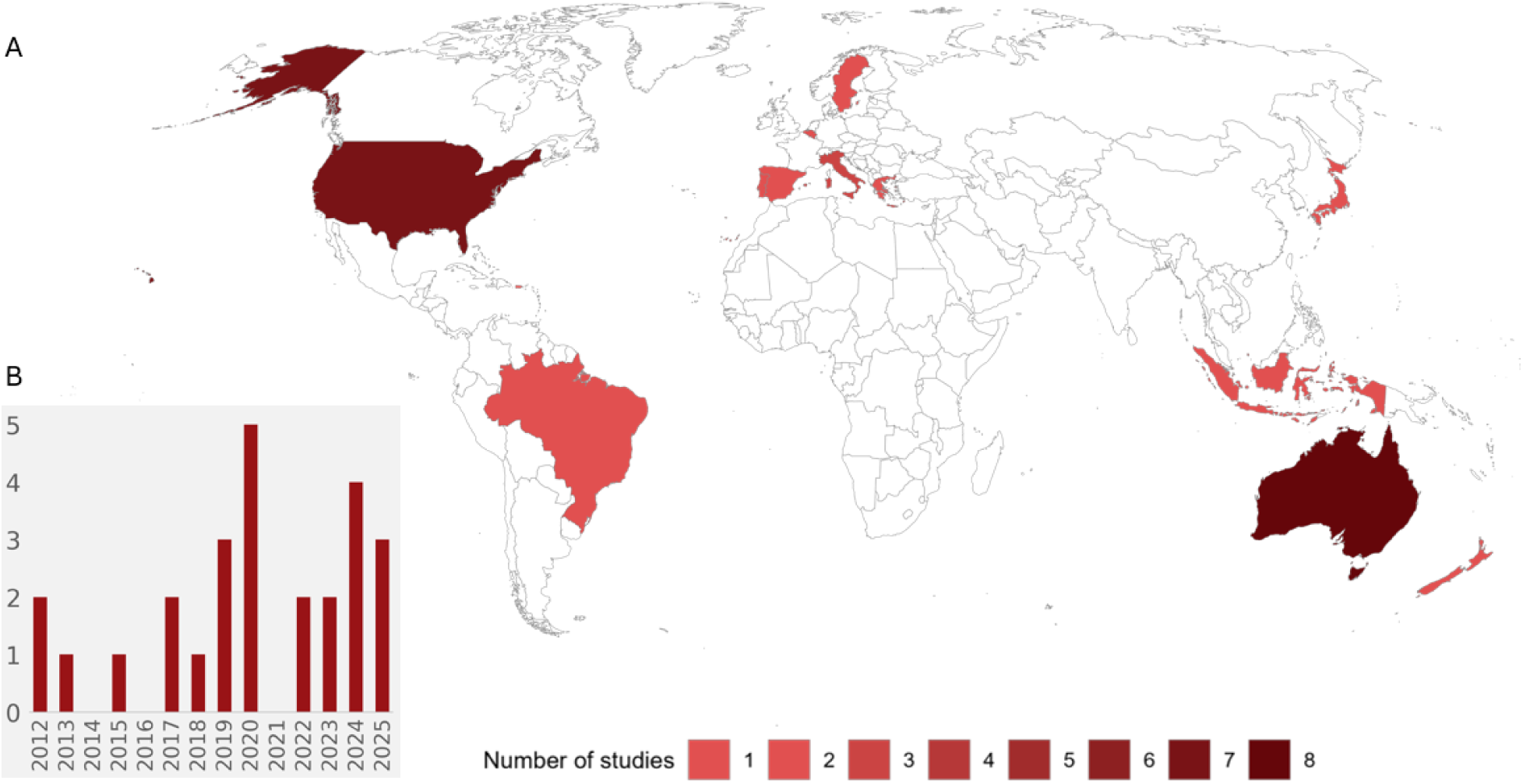
Global distribution and temporal development of studies using acoustic monitoring for the detection of invasive species. **(A)** World map showing the number of studies conducted per country. **(B)** Number of studies published per year between 2012 and 2025.

Amphibians were the most frequently studied taxonomic group in our review (11 of 26 papers), reflecting the high suitability of anurans for passive acoustic monitoring. Most studies focused on invasive frogs and toads whose mating calls are species-specific, repetitive, and often produced in conspicuous choruses, facilitating automated detection. A substantial proportion of the literature addressed the invasive cane toad (*R. marina*), and most of them in Australia (Taylor et al. 2017; Roe et al. 2018; Brodie et al. 2021; Ribeiro et al. 2022; Kimura et al. 2025; Leung et al. 2025). Beyond technical advances in automated detection, cane toad studies have also addressed broader ecological and invasion-related questions such as long-term population dynamics and their effects on native biota. Long-term acoustic monitoring along the invasion front demonstrated that cane toads rapidly became a dominant component of local anuran communities, while most native frog species persisted, albeit with altered calling intensities (Taylor et al. 2017). At larger spatial scales, passive acoustic monitoring has been integrated with occupancy modelling to assess distribution patterns and habitat associations of invasive amphibians, including *R. marina*, revealing positive links between invasion success and anthropogenically modified environments (Ribeiro et al. 2022). More recent work has focused on early detection at invasion fronts, demonstrating that deep-learning models trained in established populations in Japan can reliably identify cane toad calls in areas where the species is rare or not yet established (Kimura et al. 2025). Similarly, continent-scale classifiers based on recordings from the Australian Acoustic Observatory were able to detect cane toads across diverse environments despite regional variation in call structure (Leung et al. 2025). The study by Leung et al. (2025) furthermore showed that the classifier was able to detect cane toads reliably even in regions where it had not been trained originally, suggesting that training models with large and geographically diverse datasets helps them remain robust to differences in background noise and regional variation in mating calls.

In Australia, beyond automated sensor networks, citizen-science approaches have expanded the spatial scale of acoustic monitoring. Rowley et al. (2019) introduced FrogID, a nationwide citizen-science program in Australia that collects smartphone recordings of calling frogs, which are subsequently validated by experts. Although the project targets Australian frog biodiversity more broadly, it has proven effective in detecting invasive amphibians, including species that have established populations outside their native range. By generating geo-referenced, expert-validated acoustic records at a continental scale, FrogID demonstrates how participatory passive acoustic monitoring can contribute to early detection and range tracking of invasive amphibians (Rowley et al. 2019). Even smartphone recordings are sufficient for reliable species identification in most cases, making large-scale acoustic monitoring feasible without dedicated recording hardware (Rowley et al. 2019).

Several additional studies addressed acoustic monitoring of other invasive anuran species, including the Cuban treefrog (*Osteopilus septentrionalis*), the American bullfrog (*L. catesbeianus*), and invasive marsh frogs (*Pelophylax* spp.) in Arizona, USA, and Europe (Tennessen et al. 2013; Bisconti et al. 2019; Bota et al. 2024; Huck et al. 2024). Tennessen et al. (2013) used passive acoustic monitoring to document the presence and chorusing activity of the invasive Cuban treefrog and to assess its effects on native treefrog calling behavior. Huck et al. (2024) used passive acoustic monitoring and occupancy modeling to assess anuran distributions in stream systems in Arizona, including the invasive American bullfrog, demonstrating how passive acoustic monitoring can quantify the occurrence of invasive amphibians in relation to habitat characteristics within mixed native–invasive assemblages. In Europe, passive acoustic monitoring has been applied to track the spread of invasive water frogs, particularly the American bullfrog (*L. catesbeianus*) and Balkan marsh frogs (*Pelophylax* spp.). Bota et al. (2025) used convolutional neural networks to reliably identify American bullfrog calls in invaded habitats, showing the potential of automated classifiers for long-term monitoring of invasive amphibians in temperate European landscapes. Similarly, Bisconti et al. (2019) used passive acoustic data to detect and assess the distribution of invasive marsh frogs in Europe, whereby species-specific call characteristics can be used to distinguish expanding invasive *Pelophylax* populations from native assemblages, even within this taxonomically complex frog group.

Compared with amphibians, reptiles make far less use of acoustic communication, which may explain why our literature survey yielded only a single study addressing the potential of acoustic monitoring for the Asian house gecko (*Hemidactylus frenatus*) (Hopkins et al. 2021). Originally restricted to Southeast Asia, this species has since spread across tropical and subtropical regions worldwide, where it is classified as invasive in several areas (Weterings & Vetter 2018; Hopkins et al. 2021). Its characteristic “tik-tok” call, which gave the species its common name and results from its distinctive clucking vocalisations, is produced mainly by males during warmer months, with pronounced activity around sunset and roughly 30 minutes before sunrise (Hopkins et al. 2020). Building on this behaviour, the study by Hopkins et al. (2020) demonstrates how targeted acoustic monitoring could enable early detection of the species, helping to limit further spread.

Bird studies exclusively focused on the barred owl (*Strix varia*) in the USA, where its westward range expansion threatens the native and endangered spotted owl (*Strix occidentalis*). Since 2020, a group of scientists developed models to monitor their distribution and population growth. Lately, they managed to employ real-time monitoring of the barred owl enabling them to target and lethally remove individuals (Wood et al. 2020a, 2020b, 2024; Rugg et al. 2023; Jenkins et al. 2025).

An example of an invasive fish for which an automated classification method was developed and evaluated in Australia is the spotted tilapia (*Tilapia mariae*), which produces low frequency “knocks” or popping noises through muscular compression of its swim bladder (Kottege et al. 2012, 2015). More recently, the soniferous weakfish (*Cynoscion regalis*) became invasive on the Iberian Peninsula (since 2015) and is now threatening the native croaker (*Argyrosomus regius*) (Amorim et al. 2023). Amorim et al. (2023) developed a method to discriminate between the sounds of the invasive weakfish and those of the native fish based on pulse number and pulse rate. Furthermore, they identified the potential for using this method of passive acoustic monitoring of this invasive species in the future. Pilot studies describing the sound characteristics of the red swamp crayfish (*Procambarus clarkii*) were conducted for potential future monitoring of this species (Buscaino et al. 2012; Hisyam et al. 2020).

Invasive mammals were highly underrepresented in our literature search. Lately, a dataset of 3500 vocalizations of the common brushtail possum (*Trichosurus vulpecula*) recorded in the field was presented by McEwen et al. (2024). Based on this dataset, they developed and evaluated a model capable of automatically classifying this in New Zealand invasive opossum (McEwen et al. 2024). Similarly underrepresented are studies on soil organisms. Also soil organisms are largely underrepresented in passive acoustic monitoring. This gap is critical in contexts such as Arctic ecosystems, where invasive earthworms act as ecosystem engineers. Although direct acoustic detection of earthworms is difficult, their activity alters soil soundscapes, suggesting that indirect, large-scale monitoring may be possible through changes in belowground acoustic signatures (Keen et al. 2022).

## Passive monitoring of the invasive American bullfrog: case study

### Methods

Building on these conceptual advances and emerging applications, we further illustrate the potential of passive acoustic monitoring with a targeted case study of the invasive American bullfrog (*L. catesbeianus*). We monitored bullfrog calling activity with automated acoustic recorders (AudioMoths, Hill et al. 2018) at eleven water bodies in the Upper Rhine region near Karlsruhe in southwestern Germany, close to the major hotspot Lake Streitköpflesee in Linkenheim Hochstetten (49.1196° N, 8.3806° E; Figure 1 A+B). Simultaneous recordings over a 16-day period, from 13 June 2025 to 29 June 2025, allowed us to minimize the effects of weather fluctuations, especially temperature changes. A total of 20 AudioMoths were deployed, distributed between the study locations depending on accessibility (Figure 1A). Each AudioMoth was programmed to record 55s every minute, followed by a 5s break, from 2h before sunset until 3h after sunset at a sampling frequency of 48kHz. To automatically detect American bullfrogs in our audio recordings, we used the open-source software BirdNET Analyzer (model version V2.4, sensitivity setting of 1, minimum confidence of 0.25, and an overlap of 2 seconds; Kahl et al. 2021). BirdNET uses a convolutional neural network which is trained to identify primarily bird songs, however, also a variety of different non-bird species, including dogs, crickets, and American bullfrogs, is covered. Following Wood and Kahl (2024) we validated our bullfrog identifications by extracting 500 audio segments of 5 seconds containing bullfrog identifications, covering the whole range of confidence scores (0.25–1.00) with the implemented function *segments*. We then evaluated whether BirdNET’s identifications of American bullfrogs was correct (marked as 1) or incorrect (marked as 0) by revising the segments’ spectrograms and listening to the audio files in Kaleidoscope version 5.3.8 (Figure 3A). We used a logistic regression model to determine the BirdNET confidence score thresholds at which the probability of a correct identification was at 99% (Figure 3B) in R version 4.5.2.

**Figure 3:**
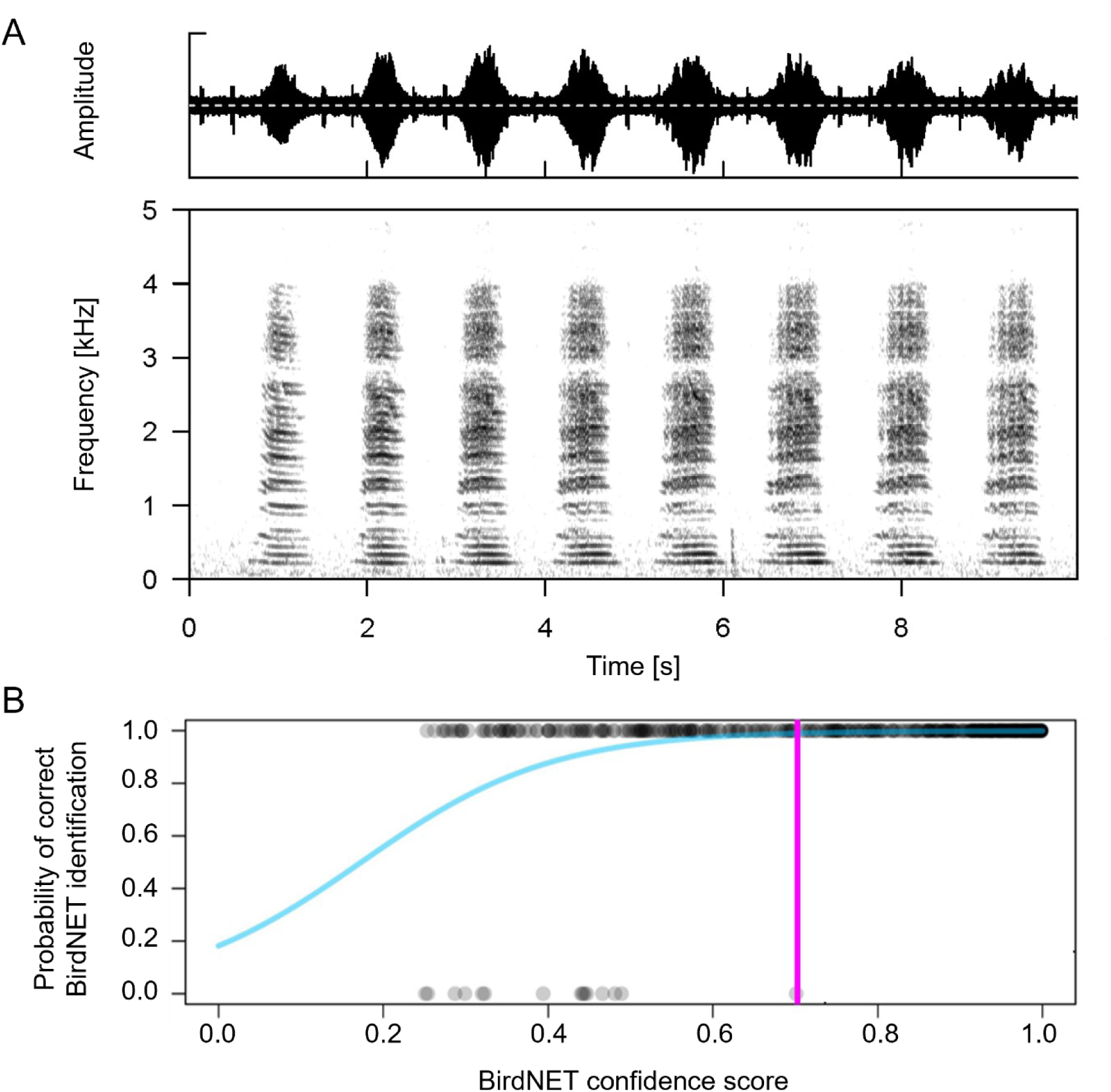
**(A)** Acoustic characteristics of an American bullfrog (*L. catesbeianus*). The amplitude plot shows changes in call intensity over time with repeated calls lasting about 1 second. The sonogram, depicting frequency (y-axis) over time (x-axis), shows the low-frequency, tonal structure of the calls. (**B)** Logistic regression showing probabilistic BirdNET score thresholds. The y-axis indicates the probability that the BirdNET prediction is correct (ranging between 0 and 1) at a certain confidence score (x-axis). Grey dots represent correct (1) and incorrect (0) BirdNET identifications. A logistic regression (blue) was used to determine the BirdNET confidence score thresholds at which the probability of a correct identification was 99%.

In addition to the AudioMoth recordings, we conducted visual shoreline surveys for bullfrog tadpoles where possible. We further compiled all available iNaturalist records of bullfrogs in the study region to assess whether passive acoustic monitoring detected bullfrog presence at sites without previously available public occurrence records.

Maps were created using R version 4.4.0 (R Core Team 2024) with packages tmap (Tennekes 2018), terra (Hijmans 2025), tmaptools (Tennekes 2025), osmdata (Padgham et al. 2017), sf (Pebesma 2018, Pebesma & Bivand 2023), rnaturalearth (Massicotte & South 2023). The background data for the world map was derived from Sentinel-2 via BKG WMS and OpenStreetMap.

## Results

In total, we recorded 1,600 hours (equivalent to over two months) of audio recordings which we analyzed with BirdNET within 24 hours. Across these recordings, BirdNET identified 56,298 snippets of 3 seconds containing bullfrog calls. Out of 500 randomly selected audio segments for subsequent validation, we classified 485 as correctly and 15 as incorrectly identified. Based on this validation, our logistic regression model indicates that BirdNET identifications of American bullfrogs with a confidence score above 0.703 had a 99% probability of being correctly identified (Figure 3B). These recordings were distributed across eight of the eleven studied water bodies (Figure 1). Additionally, we observed bullfrog tadpoles at five of these sites, which were also the only sites with available bullfrog occurrence records from citizen science in our study region (Figure 1; Supporting Information Table S2).

## Discussion

### Increasing Use of Passive Acoustic Monitoring for Invasive Animal Species

Our systematic review of the scientific literature shows that the use of passive acoustic monitoring for detecting invasive animal species is a relatively recent development which has grown rapidly over the past 14 years. The earliest studies in our dataset were published in 2012 (Buscaino et al. 2012; Kottege et al. 2012), and during the first half of the 2010s only a small number of publications explicitly addressed monitoring of invasive animal species using acoustic approaches (Kottege et al. 2012, 2015; Tennessen et al. 2013). These early studies were largely exploratory, focusing on proof-of-concept detection methods and characterization of species-specific sounds.

Starting around 2017, both the number of studies and their methodological complexity increased. In amphibians, long-term monitoring and invasion-front studies were conducted for cane toads (Taylor et al. 2017), followed by the development of automated early warning systems (Roe et al. 2018) and large-scale citizen-science initiatives (Rowley et al. 2019). In birds, a series of studies on the invasive barred owl in North America progressively advanced from distribution modelling to near real-time acoustic monitoring integrated with management actions (Wood et al. 2020a, 2020b, 2024; Rugg et al. 2023; Jenkins et al. 2025). Similar methodological shifts are evident in fish, where early automated classifiers for tilapia (Kottege et al. 2012, 2015) were followed by more recent signal discrimination frameworks for invasive weakfish in Europe (Amorim et al. 2023). The most recent studies increasingly employ deep-learning models and large, standardized datasets to enhance transferability and scalability (Bota et al. 2024; Kimura et al. 2025; Leung et al. 2025). At the same time, passive acoustic monitoring has been combined with occupancy modelling and habitat analyses to quantify the environmental factors influencing the occurrence of invasive species (Ribeiro et al. 2022; Huck et al. 2024). Emerging applications extend to mammals (McEwen et al. 2024) and even soil organisms (Keen et al. 2022), indicating a broadening taxonomic scope.

Although the total number of studies remains modest compared to more established biodiversity monitoring approaches, the clear increase of publications within the last five to ten years across multiple taxonomic groups suggests accelerating research interest and increasing methodological maturity. The field appears to be transitioning from isolated, taxon-specific case studies towards more scalable, integrative monitoring frameworks that combine automated detection, ecological modelling, and in some cases direct management intervention (Wood et al. 2024; Jenkins et al. 2025). Taken together, these developments indicate that acoustic monitoring is becoming an increasingly recognized tool in invasion ecology and biosecurity, not only for documenting presence but also for supporting early detection, tracking range expansion, and informing management responses across diverse ecosystems.

### Passive acoustic monitoring of invasive bullfrogs

Our case study demonstrates that invasive American bullfrogs (*L. catesbeianus*) can be reliably monitored using low-cost passive acoustic monitoring under field conditions in Germany. Automated pattern detection with BirdNET, combined with structured manual validation, resulted in robust species identifications of 99% classification certainty (Wood & Kahl 2024), ensuring that only highly reliable detections were considered. Subsequently, we confirmed bullfrog presence in eight of the eleven monitored lakes, including three sites for which no public occurrence records had previously been available. The detection of bullfrog calls in proximity (<100m) to a nature reserve (nature conservation area “Erlich”; protection status similar to IUCN Category IV – habitat or species management area) is particularly noteworthy, as no prior public record indicated presence of the invader. This finding suggests that current monitoring approaches may underestimate the spatial extent of invasive populations, especially in protected or less frequently surveyed areas. Given the high reproductive output and broad predatory diet of American bullfrogs (Adams & Pearl 2007; Kraus 2015), the early identification of newly colonized water bodies is ecologically significant, as it allows timely assessment of invasion risk and interventions.

Methodologically, the study shows that existing acoustic classification tools can be reliably applied to invasive amphibian monitoring when combined with structured validation. The successful use of low-cost AudioMoth recorders further shows that reliable detection is not dependent on large-scale acoustic observatories or cloud-based infrastructures (Roe et al. 2018; Leung et al. 2025). Instead, regionally deployable devices combined with clearly defined confidence thresholds can produce distribution data of sufficient certainty for applied ecological interpretation.

### Management relevance and local implementation

Invasive species management is often constrained by limited financial and personnel resources, restricted site access, and the need for repeated surveys (Haubrock et al. 2021; Dalton et al. 2024). Passive acoustic monitoring can help address some of these limitations by enabling standardized, repeated data collection with comparatively low field effort, particularly at night when invasive amphibians are most active (Dorcas et al. 2009). For management, its main value lies in confirming persistence at known sites, identifying likely breeding activity through repeated detections of mating calls, and extending surveillance to sites that are difficult to access (Gerhardt & Huber 2002; Ross et al. 2023). Rather than replacing established methods, acoustic monitoring is best used alongside visual surveys, targeted field checks, and citizen science, where it can help prioritize follow-up actions and support more efficient allocation of limited resources (Sugai et al. 2019; Hoefer et al. 2026).

### Transferability to other invasive species and contexts

Globally, many invasive taxa produce frequent and distinctive sounds, including other amphibians, birds, reptiles and even mammals as shown in our literature review, making them suitable candidates for acoustic monitoring (Taylor et al. 2017; Roe et al. 2018; Sugai et al. 2019; Bota et al. 2024; McEwen et al. 2024; Wood et al. 2024; Jenkins et al. 2025; Leung et al. 2025). More generally, passive acoustic monitoring is particularly useful for species that produce identifiable sounds consistently enough to allow reliable detection over time and across diverse habitats (Sugai et al. 2019; Ross et al. 2023). Importantly, the studies included in our review are likely to represent only a fraction of the potential applications of passive acoustic monitoring of invasive species. Beyond the taxa currently investigated, growing interest exists in applying acoustic methods to flying insects through their flight sounds (Fengler et al. 2026). Wingbeat frequencies have been described as species-specific “acoustic fingerprints” for some invertebrates, suggesting that passive acoustic monitoring could even support detection of invasive insect species in the future (Mankin et al. 2011; Kawakita & Ichikawa 2019; Hall et al. 2025). For example, acoustic monitoring systems deployed near honeybee hives could enable early detection of the invasive Asian hornet (*Vespa velutina*) as shown by Hall et al. (2025). Coupled with emerging real-time acoustic monitoring technologies, such approaches may provide opportunities for continuous monitoring and rapid management responses to the presence of invasive species (Wood et al. 2024). Commercial systems such as ecoPi (OekoFor GbR, Freiburg, Germany) or BirdWeather PUC (Scribe Labs Inc., California, USA), as well as custom Raspberry Pi-based setups (Raspberry Pi Ltd., Cambridge, UK), can process audio in the field, detect target species automatically, and transmit alerts via cellular networks. Although still a recent and sparsely documented area, these methods have strong potential to enable earlier interventions in invasion ecology.

## Conclusion

Our systematic literature review and case study together demonstrate that acoustic monitoring of animal species is becoming an increasingly useful tool in invasion ecology. Its value lies not only in repeated and cost-effective monitoring over broad temporal and spatial scales, but also in its compatibility with ongoing advances in automated species identification, in-field data processing, and emerging real-time monitoring workflows. When implemented with transparent validation, clear monitoring protocols, and integration with existing survey methods, passive acoustic monitoring has strong potential to support early intervention and to improve the allocation of limited management resources.

## CRediT authorship contribution statement

**David Becker**: Data collection, Methodology, Visualisation, Writing – original draft. **Marit K. Kasten**: Data analysis, Methodology, Visualisation, Writing – original draft. **Torben Weber**: Data collection, Methodology, Visualisation, Writing – original draft. **Ingo Grass**: Writing – review & editing. **Thomas Hiller**: Methodology, Supervision, Visualisation, Writing – original draft.

