## Supporting Information for "Passive acoustic monitoring as a tool for early detection of invasive animal species: a systematic review and a case study"

Table S1. Studies included in the systematic literature review on the use of passive acoustic monitoring for invasive animal species, including publication details, study region, taxonomic group, and target species.

| **Author** | **Year** | **Title** | **Country** | **Group** | **Species** |
|---|---|---|---|---|---|
| Kimura et al. | 2025 | Deep learning-based detector of invasive alien frogs, *Polypedates leucomystax* and *Rhinella marina*, on an island at invasion front | Japan | Amphibians | Common tree frog (*Polypedates leucomystax*) and Cane toad (*Rhinella marina*) |
| Leung et al. | 2025 | Advancing invasive species monitoring: A free tool for detecting invasive cane toads using continental-scale data | Australia | Amphibians | Cane toad (*Rhinella marina*) |
| Huck et al. | 2024 | Anuran occupancy varies with stream characteristics and flow across Arizona wilderness areas | Arizona | Amphibians | Frogs including invasive species |
| Bota et al. | 2024 | Passive acoustic monitoring and automated detection of the American bullfrog | Belgium, Italy, Spain | Amphibians | American bullfrog (*Lithobates catesbeianus*) |
| Ribeiro et al. | 2022 | Passive Acoustic Monitoring as a Tool to Investigate the Spatial Distribution of Invasive Alien Species | USA | Multiple species | Birds, mammals, frogs |
| Brodie et al | 2020 | Acoustic monitoring reveals year-round calling by invasive toads in tropical Australia | Australia | Amphibians | Cane toad (*Rhinella marina*) |
| Bisconti et al. | 2019 | Balkan marsh frogs *Pelophylax kurtmuelleri* (Gayda, 1940) introduced in the Aspromonte National Park, southern Italy | Italy | Amphibians | The balkan frog (*Pelophylax kurtmuelleri*) |
| Rowley et al. | 2019 | FrogID: Citizen scientists provide validated biodiversity data on frogs of Australia | Australia | Amphibians | Frogs including invasive species |

| | | | | | |
|---|---|---|---|---|---|
| Roe et al. | 2018 | Catching Toad Calls in the Cloud: Commodity Edge Computing for Flexible Analysis of Big Sound Data | Australia | Amphibians | Cane toad (*Rhinella marina*) |
| Taylor et al. | 2017 | Impact of cane toads on a community of Australian native frogs, determined by 10 years of automated identification and logging of calling behaviour | Australia | Amphibians | Cane toad (*Rhinella marina*) |
| Tennessen et al. | 2013 | Impacts of acoustic competition between invasive Cuban treefrogs and native treefrogs in southern Florida | USA | Amphibians | Cuban treefrog (*Osteopilus septentrionalis*) |
| Wood et al. | 2020 | Early detection of rapid Barred Owl population growth within the range of the California Spotted Owl advises the Precautionary Principle | USA | Birds | Barred Owl (*Strix varia*) |
| Wood et al. | 2020 | Using the ecological significance of animal vocalizations to improve inference in acoustic monitoring programs | USA | Birds | Barred Owl (*Strix varia*) |
| Rugg et al. | 2023 | Western screech-owl occupancy in the face of an invasive predator | USA | Birds | Barred Owl (*Strix varia*) |
| Wood et al. | 2024 | Real-time acoustic monitoring facilitates the proactive management of biological invasions | USA | Birds | Barred Owl (*Strix varia*) |
| Jenkins et al. | 2025 | Congener Conflict: Invasive Barred Owl Landscape Use Across the Range of the Northern Spotted Owl | USA | Birds | Barred Owl (*Strix varia*) |
| Kottege et al. | 2012 | Classification of Underwater Broadband Bio-acoustics Using Spectro-Temporal Features | Australia | Fish | Spotted tilapia (*Tilapia mariae*) |
| Kottege et al. | 2015 | Automated detection of broadband clicks of freshwater fish using spectro-temporal features | Australia | Fish | Spotted tilapia (*Tilapia mariae*) |

| | | | | | |
|---|---|---|---|---|---|
| Demertzis et al. | 2017 | A deep spiking machine-hearing system for the case of invasive fish species | Mediterranean Sea | Fish | Native vs invasive species |
| Raick et al. | 2020 | Yellow-eyed piranhas produce louder sounds than red-eyed piranhas in an invasive population of *Serrasalmus marginatus* | Brazil | Fish | Spotted piranha (*Serrasalmus marginatus*) |
| Amorim et al. | 2023 | Detection of invasive fish species with passive acoustics: Discriminating between native and non-indigenous sciaenids | Portugal | Fish | Soniferous weakfish (*Cynoscion regalis*) |
| McEwen et al. | 2024 | An invasive species model and dataset for bioacoustic monitoring of common brushtail possum | New Zealand | Mammals | Common brushtail possum (*Trichosurus vulpecula*) |
| Buscaino et al. | 2012 | The underwater acoustic activities of the red swamp crayfish *Procambarus clarkii* | Italy | Decapoda | Red swamp crayfish (*Procambarus clarkii*) |
| Hisyam et al. | 2020 | Sound characteristic of *Procambarus clarkii* | Indonesia | Decapoda | Red swamp crayfish (*Procambarus clarkii*) |
| Keen et al. | 2022 | Non-native species change the tune of tundra soils: Novel access to soundscapes of the Arctic earthworm invasion | Sweden | Arthropods | Earthworm |
| Hopkins et al. | 2019 | Calling behaviour in the invasive Asian house gecko (*Hemidactylus frenatus*) and implications for early detection | Australia | Reptiles | Asian house gecko (*Hemidactylus frenatus*) |

Table S2. Summary of AudioMoth locations, acoustic detections of American bullfrogs, field observations of tadpoles/toads, and associated iNaturalist records for the monitored water bodies.

| Audio Moth ID | Coord. | Audio detected | Filtered 99perc BirdNET identifications | Bullfrog tadpoles/ toads | Inat. detected | Inat. detected number |
|---|---|---|---|---|---|---|
| 17 | 49.120510, 8.377198 | yes | 5977 | yes | yes | 29 |
| 36 | 49.122501, 8.377053 | yes | 29656 | yes | yes | 29 |
| 59 | 49.120342, 8.381930 | yes | 19 | yes | yes | 29 |
| 74 | 49.116772, 8.381233 | no | 0 | yes | yes | 29 |
| 64 | 49.126858, 8.389052 | yes | 2349 | no | no | 0 |
| 70 | 49.124497, 8.398581 | yes | 1192 | no | no | 0 |
| 24 | 49.156124, 8.393748 | no | 0 | no | no | 0 |
| 51 | 49.160980, 8.396600 | no | 0 | no | no | 0 |
| 41 | 49.154846, 8.385703 | yes | 15 | yes | yes | 1 |
| 50 | 49.152805, 8.386866 | yes | 1 | yes | yes | 1 |
| 62 | 49.191212, 8.468065 | no | 0 | no | no | 0 |
| 66 | 49.188179, 8.457278 | yes | 5 | no | no | 0 |
| 53 | 49.185108, 8.463417 | no | 0 | no | no | 0 |
| 56 | 49.104214, 8.389495 | yes | 29 | no | yes | 1 |
| 68 | 49.108547, 8.396985 | yes | 56 | no | yes | 1 |
| 73 | 49.114258, 8.390585 | yes | 77 | no | yes | 1 |
| 57 | 49.083855, 8.371782 | yes | 1 | yes | yes | 1 |
| 58 | 49.0881894, 8.375050 | yes | 2 | yes | yes | 26 |

| | | | | | | |
|---|---|---|---|---|---|---|
| 71 | 49.081070, 8.368320 | no | 0 | yes | yes | 1 |
| 54 | 49.067062, 8.359282 | yes | 2 | no | no | 0 |
